# Different excitation-inhibition correlations between spontaneous and tone-evoked activity in primary auditory cortex neurons

**DOI:** 10.1101/2021.12.20.473450

**Authors:** Katherine C. M. Chew, Vineet Kumar, Andrew Y. Y. Tan

**Affiliations:** Department of Physiology, Yong Loo Lin School of Medicine, National University of Singapore Healthy Longevity Translational Research Programme, Yong Loo Lin School of Medicine, National University of Singapore; Cardiovascular Translational Research Programme, Yong Loo Lin School of Medicine, National University of Singapore; Neurobiology Programme, Life Sciences Institute, National University of Singapore, 28 Medical Drive, Singapore 117456, Republic of Singapore

**Keywords:** primary auditory cortex, spontaneous activity, tone-evoked responses, synaptic excitation, synaptic inhibition, correlation, in vivo whole-cell

## Abstract

Tone-evoked synaptic excitation and inhibition are highly correlated in many neurons with V-shaped tuning curves in the primary auditory cortex of pentobarbital-anesthetized rats. In contrast, there is less correlation between spontaneous excitation and inhibition in visual cortex neurons under the same anesthetic conditions. However, it was not known whether the primary auditory cortex resembles visual cortex in having spontaneous excitation and inhibition that is less correlated than tone-evoked excitation and inhibition. Here we report whole-cell voltage-clamp measurements of spontaneous excitation and inhibition in primary auditory cortex neurons of pentobarbital-anesthetized rats. The larger excursions of both spontaneous excitatory and inhibitory currents appeared to consist of distinct events, with the inhibitory event rate typically lower than the excitatory event rate. We use the ratio of the excitatory event rate to the inhibitory event rate, and the assumption that the excitatory and inhibitory synaptic currents can each be reasonably described as a filtered Poisson process, to estimate the maximum spontaneous excitatory-inhibitory correlation for each neuron. In a subset of neurons, we also measured tone-evoked excitation and inhibition. In neurons with V-shaped tuning curves, although tone-evoked excitation and inhibition were highly correlated, the spontaneous inhibitory event rate was typically sufficiently lower than the spontaneous excitatory event rate to indicate a lower excitatory-inhibitory correlation for spontaneous activity than for tone-evoked responses.

## Introduction

Primary auditory cortex neurons typically spike in response to a narrow range of tone frequencies at low tone intensities, and to an increasingly broader range of frequencies as intensity is increased, which results in V-shaped tuning curves. In anesthetized and in awake animals, the underlying tone-evoked synaptic excitation and inhibition in such neurons also have V-shaped tuning curves, with the magnitude of excitation and inhibition typically highly correlated as tone frequency and intensity are varied, except for neurons in layer 6 (Zhou et al., 2010). Across neurons in layers 2/3 to 5, the correlation (Pearson correlation coefficient) between tone-evoked excitation and inhibition ranges from approximately 0.6 to 0.9, with an average value of approximately 0.8 (Wehr and Zador, 2003; Zhang et al., 2003; Tan and Wehr, 2009; Zhou et al., 2014).

In contrast, spontaneous excitation and inhibition in visual cortex neurons of pentobarbital-anesthetized rats are more weakly correlated. Spontaneous excitatory and inhibitory currents appeared to consist of distinct events. If excitation and inhibition were perfectly correlated, excitatory and inhibitory event rates would be equal. However, the inhibitory event rate was typically lower than the excitatory event rate, such that the average correlation between spontaneous excitation and inhibition was less than 0.6 (Tan et al., 2013).

However, the correlation between spontaneous excitation and inhibition in the primary auditory cortex has not been previously reported. One possibility is that spontaneous excitation and inhibition in the primary auditory cortex resembles that in the visual cortex in being less correlated than tone-evoked excitation and inhibition. Alternatively, the correlation between spontaneous excitation and inhibition may be as high as that for tone-evoked excitation and inhibition, which may be the case if spontaneous activity resembles activity evoked by random tone stimulation (DeWeese and Zador, 2006).

Accordingly, we carried out whole-cell voltage clamp measurements of spontaneous excitation and inhibition in primary auditory cortex neurons of pentobarbital-anesthetized rats. We found that the larger excursions of spontaneous excitatory and inhibitory currents appeared to consist of distinct events, with the inhibitory event rate typically lower than the excitatory event rate. We use the ratio of the excitatory event rate to the inhibitory event rate, and the assumption that the excitatory and inhibitory synaptic currents can each be reasonably described as a filtered Poisson process, to estimate the maximum spontaneous excitatory-inhibitory correlation for each neuron. In a subset of neurons, we also measured tone-evoked excitation and inhibition. In neurons with V-shaped tuning curves, although tone-evoked excitation and inhibition were highly correlated, the spontaneous inhibitory event rate was typically sufficiently lower than the spontaneous excitatory event rate to indicate that spontaneous excitation and inhibition are less correlated than tone-evoked excitation and inhibition.

## Experimental procedures

### Physiology

All experiments in this study were approved by the National University of Singapore Institutional Animal Care and Use Committee, and were in accordance with the U.S. National Institutes of Health Guide for the Care and Use of Laboratory Animals (NIH Publications No. 80-23, revised 1996). Experiments were carried out in a sound-attenuating chamber. We used female Sprague-Dawley rats about 2-3 months old, weighing 200-300 g. Each rat was anesthetized by intraperitoneal injection of sodium pentobarbital (50-80 mg/kg), with the dose adjusted to make the rat areflexic. The rat was maintained in an areflexic state for the rest of the experiment by further intraperitoneal injections of sodium pentobarbital (20-60 mg/kg) when necessary. The rat was placed on a heating pad, and its temperature was maintained at ∼37°C. Prior to any skin incision, lidocaine was injected subcutaneously at the incision site. A tracheotomy was performed to secure the airway. The head was held fixed by a custom-made device that clamped it at both orbits and the palate, leaving the ears unobstructed. A cisternal drain was performed. The right auditory cortex was exposed by retracting the skin and muscle overlying it, followed by a craniotomy and a durotomy. The cortical surface was kept moist with normal saline or 4% agarose in normal saline. The location of primary auditory cortex was determined by coarse mapping of multiunit spike responses at 500-600 μm below the pial surface with a parylene-coated tungsten electrode (MicroProbes for Life Sciences, Gaithersburg, MD, USA).

### Whole cell recordings

Blind whole-cell recordings in vivo were obtained (Pei et al., 1991; Ferster and Jagadeesh, 1992; Tan et al., 2004; Tan et al., 2013). A silver wire, one end of which was coated with silver chloride, served as the reference electrode against which potentials were measured; its chlorided end was inserted into muscle near the base of the skull. The reference electrode was assigned a potential of 0 mV. The potential of the cerebrospinal fluid was assumed to be uniform and equal to that of the reference electrode. Pipettes (3–6 MΩ) were pulled from 1.2 mm outer diameter, 0.7 mm inner diameter KG-33 or 7740 borosilicate glass capillaries (King Precision Glass, Claremont, CA, USA) on a P-2000 micropipette puller (Sutter Instruments, Novato, CA, USA) to record from neurons 400 to 950 μm below the cortical surface, corresponding to positions in layers 2/3 to 5 (Kaur et al., 2005; Sakata and Harris, 2009). Pipettes were filled with solution A, solution B or solution C; there were no apparent differences in results among the solutions, so we have combined data from all solutions for our analyses. Solution A contained (in mm) 135 Cs-methanesulfonate, 5 TEA-Cl, 5 QX-314, 0.5 EGTA, 10 HEPES, 10 phosphocreatine disodium, pH 7.3; solution B contained 135 Cs-methanesulfonate, 5 TEA-Cl, 5 QX-314, 4 BAPTA, 10 HEPES, 10 phosphocreatine disodium, pH 7.3; solution C contained 135 Cs-methanesulfonate, 5 TEA-Cl, 5 QX-314, 0.5 MK-801, 4 BAPTA, 10 HEPES, 10 phosphocreatine disodium, pH 7.3; pH was adjusted with CsOH (Sigma-Aldrich, St. Louis, MO, USA). Voltage clamping was performed with a MultiClamp 700B patch-clamp amplifier (Molecular Devices, Sunnyvale, CA, USA). Series resistances ranged from 20 to 80 MΩ, with a median value of 45 MΩ. Excitatory synaptic currents were recorded by voltage clamping with a command potential of -85 mV; after accounting for series resistance and subtraction of an approximately 10 mV junction potential, the median membrane potential for recording spontaneous excitation was approximately -85 mV, near the calculated inhibitory reversal potential of -90 mV. Inhibitory synaptic currents were recorded by voltage clamping with a command potential of 10 mV or 20 mV; after accounting for series resistance and subtraction of an approximately 10 mV junction potential, the median membrane potential for recording spontaneous inhibition was approximately -10 mV, near the calculated excitatory reversal potential of 0 mV. Traces were low-pass filtered at 4000 Hz, and digitized at 8064.52 Hz.

### Stimuli

Stimuli were delivered by a calibrated free field speaker (Peerless by Tymphany XT25TG30-04) directed toward the left ear. Stimuli consisted of 350 pure tone pips, each with one of 20 frequencies spanning 0.5-40.317 kHz in 1/3 octave steps, and 7 intensities spanning 10-70 dB SPL in 10 dB steps, each with a 30-ms duration and 3-ms linear rising and falling phases. The inter-tone interval was 750 ms.

### Analysis of spontaneous activity

To detect events in spontaneous excitatory and inhibitory currents by their peaks, we subtracted baselines from traces, and used the MATLAB function smooth to implement lowess smoothing with a 20-ms span to avoid spurious peaks due to high-frequency noise and voltage-clamp errors. We then used the MATLAB function findpeaks to detect events with a 20-ms minimum width and peaks with amplitudes greater than 0.1 of the maximum amplitude.

Confidence intervals (95%) for event rates were estimated as Rao score confidence intervals; confidence intervals (95%) for event rate ratios were estimated as MOVER Rao score confidence interval based on the Fieller’s theorem (Li et al., 2014). The values of the confidence limits (95%) for the maximum possible spontaneous excitatory-inhibitory correlation were estimated as those corresponding to the values of the confidence limits (95%) for the ratio of the excitatory event rate to the inhibitory event rate.

### Analysis of tone-evoked responses

We defined tuning analysis regions for analyzing tone-evoked responses. A tuning analysis region contained (i) all tones within the tuning curve, (ii) tones up to 1/6 octaves below and above the lower or upper frequency boundaries at each intensity of the tuning curve, and (iii) all tones at 10 dB SPL, which was typically below the intensity threshold for tone-evoked responses. The tuning curve of a neuron was defined to contain all tones for which the current within a defined time window after tone onset exceeded 3 standard deviations of the currents 50 ms before tone onset, and which were one of at least 3 successive frequencies for which the currents exceeded the 3 standard deviation criterion. For excitation, the defined time window was 10-30 ms after tone onset; for inhibition, it was 10-50 ms after tone onset. For computing the correlation between tone-evoked excitation and inhibition for a neuron, as well as for computing the ratio of the number of tone-evoked inhibitory responses to the number of tone-evoked excitatory responses, analysis for each neuron was restricted to tones within the combined tuning analysis regions for excitation and inhibition.

In some neurons, for which the tone stimulus set was presented twice (before and after recording spontaneous excitation and inhibition), the correlation between tone-evoked excitation and inhibition was calculated over responses from both presentations of the tone stimulus set. Confidence intervals (95%) for the excitatory-inhibitory correlation for tone-evoked responses were calculated using the MATLAB function corrcoeff.

## Results

We performed whole-cell voltage-clamp recordings of spontaneous excitatory and inhibitory synaptic currents in 70 primary auditory cortex neurons of pentobarbital-anesthetized rats. We used a cesium-based intracellular solution containing pharmacological blockers of voltage-dependent channels to ensure that recorded currents were synaptic with minimal contributions from intrinsic channels. We recorded excitatory synaptic currents by clamping the membrane voltage near the inhibitory reversal potential, and inhibitory synaptic currents by clamping near the excitatory reversal potential.

For both spontaneous excitatory and inhibitory currents, fluctuations larger than 0.1 of the maximum amplitude consisted of distinct events. We identified events larger than 0.1 of the maximum amplitude by their peaks. As shown in two example neurons (Fig. 1A, B), the excitatory event rate was typically greater than the inhibitory event rate. Across neurons, the excitatory event rate had a mean value of 3.0 per second and a median value of 3.2 s^-1^; the inhibitory event rate had a mean value of 1.2 s^-1^and a median value of 1.0 s^-1^; the median excitatory event rate was greater than the median inhibitory event rate (Wilcoxon sign rank test, p = 4.8 × 10^−13^; Fig. 2A). Across neurons, the ratio of the excitatory event rate to the inhibitory event rate in each neuron (EI rate ratio) had a mean value of 3.5 and a median value of 2.6. These mean and median EI rate ratios for the primary auditory cortex were similar to the mean EI rate ratio of 2.7 previously reported for spontaneous excitation and inhibition in the visual cortex (Tan et al., 2013).

**Fig. 1.**
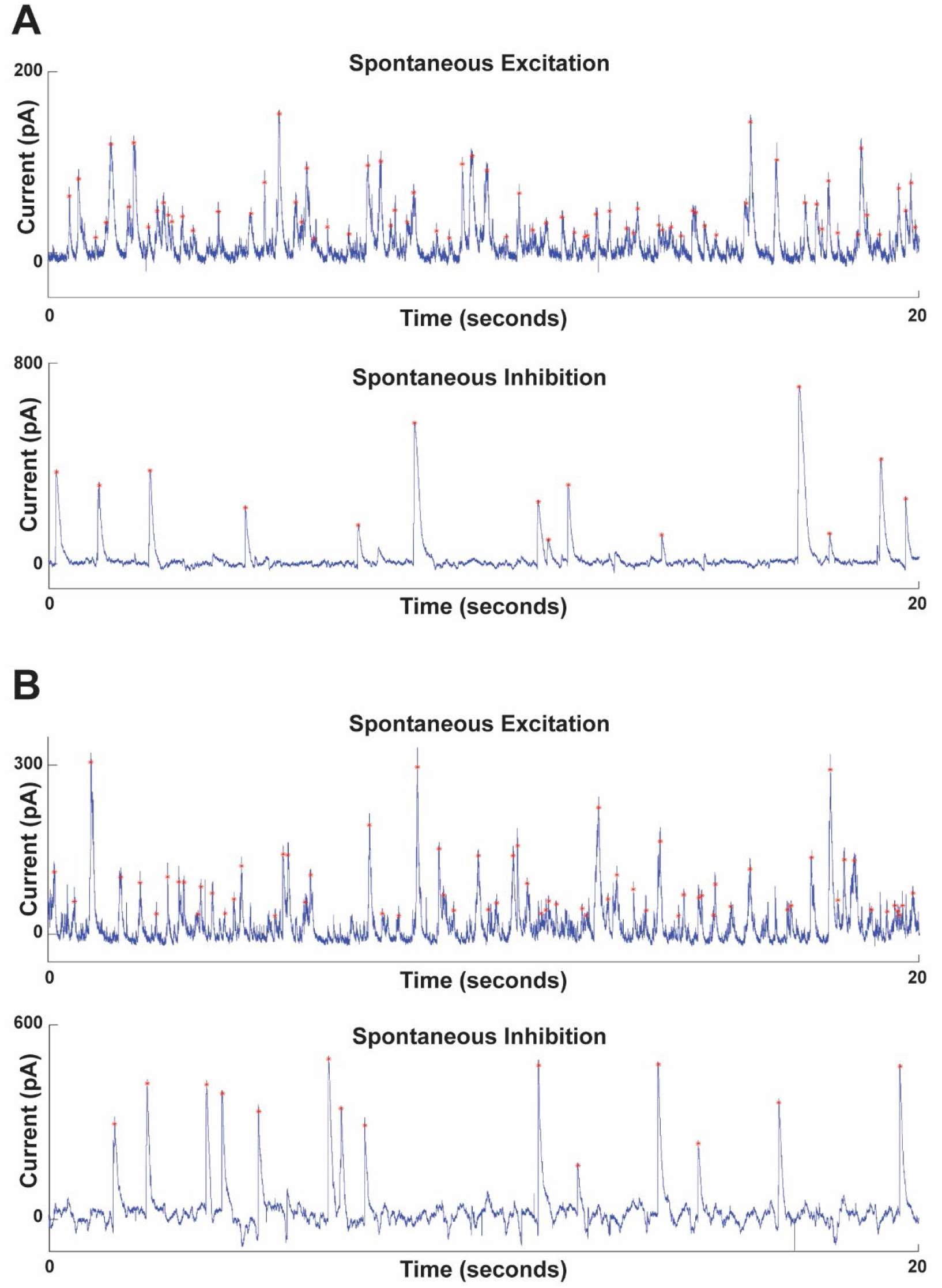
Records of spontaneous synaptic excitation and inhibition from two example neurons. Each of (A) and (B) show records from a different neuron. For each neuron, the top panel shows spontaneous excitatory synaptic currents; the bottom panel shows spontaneous inhibitory synaptic currents; in the top panels, upward deflections represent inward current; in the bottom panels, downward deflections represent inward currents. Events detected by their peaks are marked with red asterisks.

**Fig. 2.**
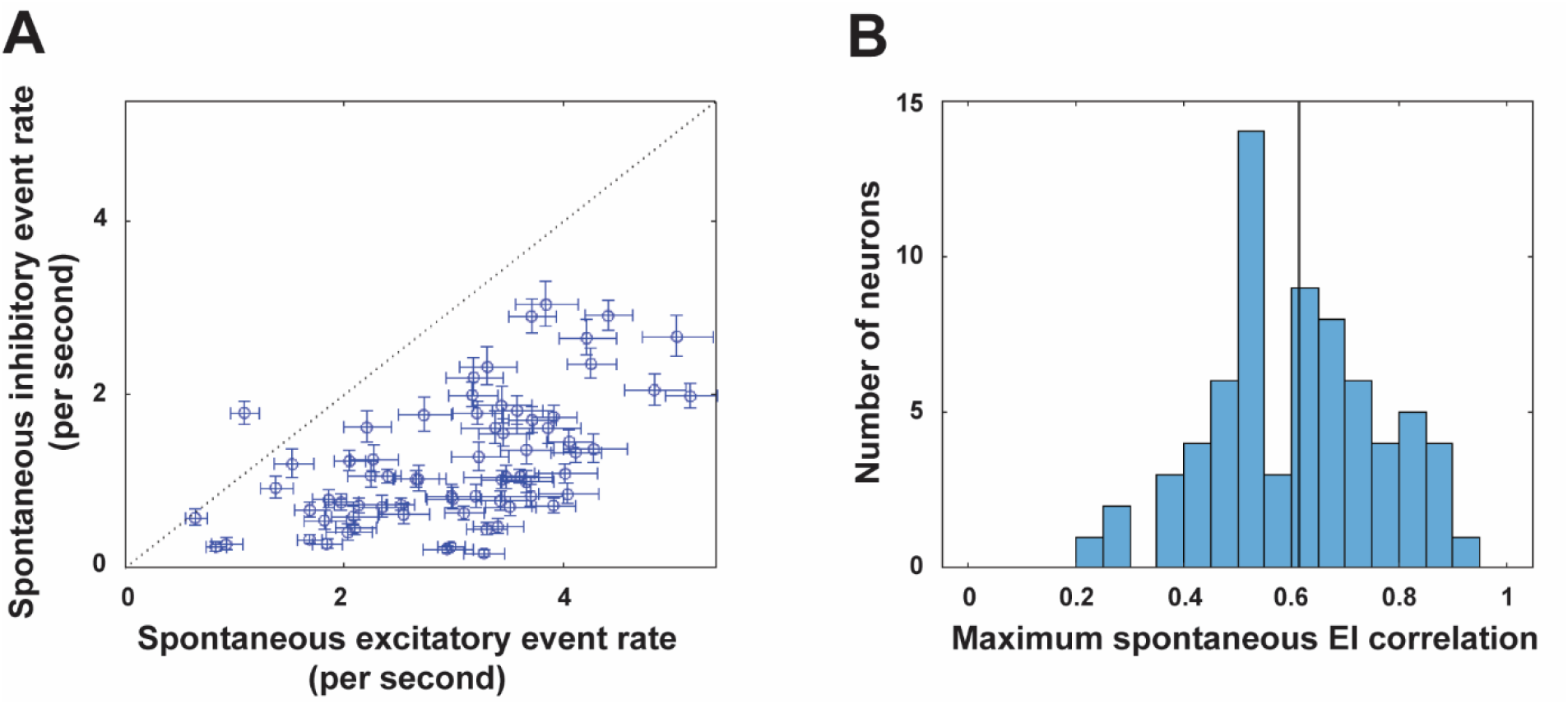
Spontaneous excitatory event rates are greater than inhibitory event rates. Data are shown for 70 neurons from which spontaneous excitatory and inhibitory currents were recorded. (A) Spontaneous inhibitory event rate versus spontaneous excitatory event rate. Error bars represent 95% confidence intervals. (B) Histogram of maximum spontaneous EI correlations estimated from spontaneous EI rate ratios.

If spontaneous excitation and inhibition were perfectly correlated, excitatory and inhibitory event rates would be identical, and the EI rate ratio would be 1. Hence, an EI rate ratio that is different from 1 implies that excitation and inhibition are less than perfectly correlated. It was previously shown that the maximum possible spontaneous excitatory-inhibitory correlation (maximum spontaneous EI correlation) *r*_*max*_ for an EI rate ratio *x* is given by *r*_*max*_ = *x*^−1/2^, if spontaneous excitatory and inhibitory currents can each be modelled as a filtered Poisson process (Tan et al., 2013). For a filtered Poisson process, the distribution of inter-event intervals is exponential, or equivalently, the logarithm of the distribution of inter-event intervals is linear. Across our synaptic current records, the linear correlation coefficient for the logarithm of the distribution of inter-event intervals was typically greater than 0.9 if the record was long enough to contain at least 250 events; the linear correlation coefficient tended to increase with the number of events recorded, suggesting that most linear correlation coefficients with smaller values were artifacts of shorter record durations, rather than indicating synaptic currents that could not be reasonably described as filtered Poisson processes. We have thus assumed that excitatory and inhibitory synaptic currents in the primary auditory cortex are reasonably described as filtered Poisson processes, and computed the maximum spontaneous EI correlation for each neuron given its EI rate ratio (Fig. 2B). Across neurons, the maximum spontaneous EI correlation ranged from 0.23 to 0.95, and had a mean value of 0.61 and a median value of 0.61. These average maximum spontaneous EI correlation values for the primary auditory cortex are similar to the average value of 0.61 reported for the visual cortex (Tan et al., 2013).

These data (Fig. 1, 2) show that spontaneous excitation and inhibition in the primary auditory cortex resembles that in the visual cortex, and suggests that spontaneous excitation and inhibition in the primary auditory cortex are less correlated than tone-evoked excitation and inhibition. To investigate whether this holds even in neurons with V-shaped tuning curves and highly correlated tone-evoked excitation and inhibition, in some neurons we recorded tone-evoked excitation and inhibition in addition to spontaneous activity.

We restrict all following analyses to 40 neurons whose excitatory tuning curves were clearly V-shaped by visual inspection. Fig. 1A & B showed spontaneous excitation and inhibition records from 2 example neurons. Fig. 3A & B show the corresponding tone-evoked excitation and inhibition for each of those 2 example neurons. In each example neuron, tone-evoked excitation and inhibition both exhibited V-shaped tuning curves, with the magnitude of excitation and inhibition within a defined time window highly correlated across the frequency and intensity of tones within the combined excitatory and inhibitory tuning analysis regions. For excitation, the defined time window was 10-30 ms after tone onset; for inhibition, it was 10-50 ms after tone onset. Across our sample of neurons with V-shaped tuning curves, the correlation between tone-evoked excitation and inhibition (tone-evoked EI correlation) ranged from 0.55 to 0.91, with a mean value of 0.75 and a median value of 0.77. This was consistent with previous reports that the tone-evoked EI correlation ranges from approximately 0.6 to 0.9 (Wehr and Zador, 2003; Zhang et al., 2003; Tan and Wehr, 2009; Zhou et al., 2014), with an average value of approximately 0.8 (Wehr and Zador, 2003).

**Fig. 3.**
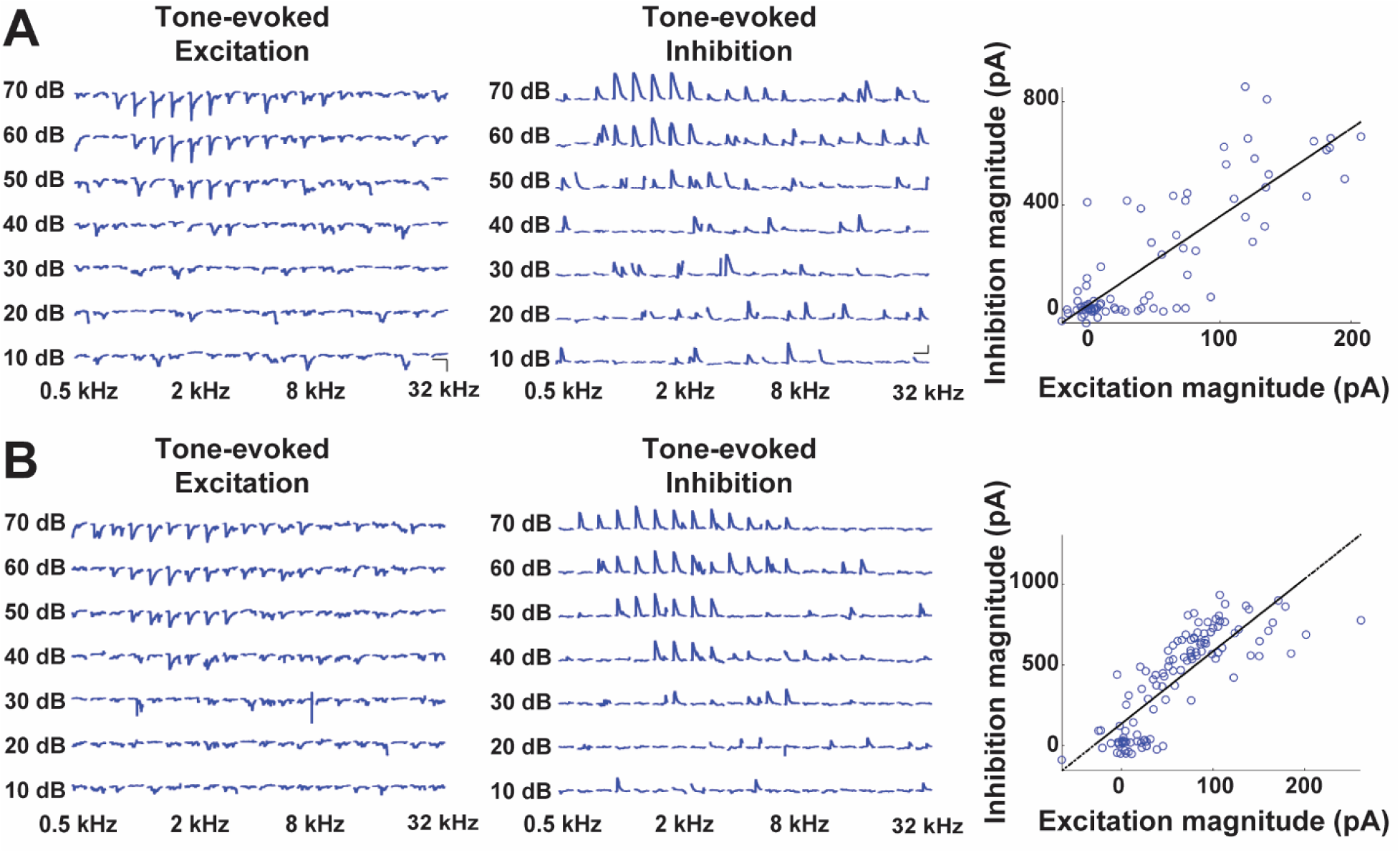
Records of spontaneous synaptic excitation and inhibition from two example neurons. Each of (A) and (B) show records from a different neuron, and are respectively from the same neuron as in Fig. 1A and Fig. 1B. For each neuron, the left panel shows tone-evoked excitatory synaptic currents for tones of various frequencies and intensities; the middle panel shows tone-evoked inhibitory synaptic currents; the right panel shows inhibitory synaptic current magnitude versus excitatory current magnitude. In the left and middle panels, downward deflections represent inward current.

For spontaneous activity across the sample of neurons with V-shaped tuning curves, the excitatory event rate had a mean value of 3.1 per second and a median value of 3.2 s^-1^; the inhibitory event rate had a mean value of 1.3 s^-1^ and a median value of 1.1 s^-1^; the median excitatory event rate was greater than the median inhibitory event rate (Wilcoxon sign rank test, p = 4.8 × 10^−8^; Fig. 4A). The EI rate ratio had a mean value of 3.4 and a median value of 2.6. The maximum spontaneous EI correlation ranged from 0.23 to 0.89, and had a mean value of 0.62 and a median value of 0.62 (Fig. 4B). Spontaneous activity in the sample of neurons with V-shaped tuning curves was thus comparable to that in the total sample of neurons with spontaneous activity records (Fig. 2).

**Fig. 4.**
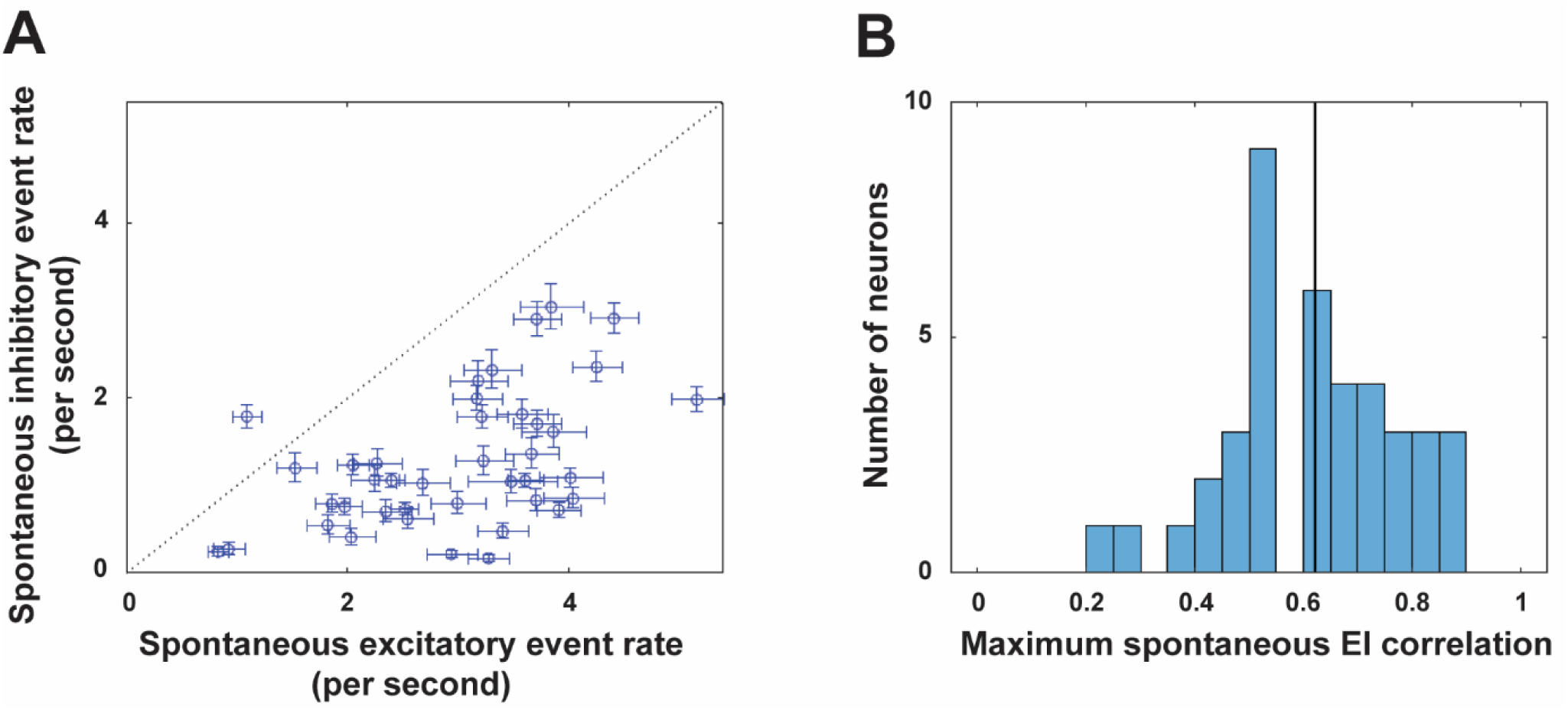
Spontaneous excitatory event rates are greater than inhibitory event rates. Data are shown for the subset of 40 neurons (out of the 70 shown in Fig. 2) from which both spontaneous and tone-evoked excitatory and inhibitory currents were recorded. (A) Spontaneous inhibitory event rate versus spontaneous excitatory event rate. Error bars represent 95% confidence intervals. (B) Histogram of maximum spontaneous EI correlations estimated from spontaneous EI rate ratios.

For each neuron in the sample of neurons with V-shaped tuning curves, we now compare spontaneous excitation and inhibition with tone-evoked excitation and inhibition. First, we compute the ratio of the number of tone-evoked inhibitory events to the number of tone-evoked excitatory events (tone-evoked I/E rate ratio) for tones within the combined excitatory and inhibitory tuning analysis regions. For each tone, we considered there to be an excitatory event if the excitatory current 10-30 ms after tone onset was larger than 0.1 of the maximum amplitude in the spontaneous excitatory current record for that neuron; we considered there to be an inhibitory event if the inhibitory current 10-50 ms after tone-onset was larger than 0.1 of the maximum amplitude in the spontaneous inhibitory current record for that neuron. Across neurons the tone-evoked I/E rate ratio had mean 0.93 and median 0.96, which being close to 1, is consistent with almost every tone that evoked an excitatory event also evoking an inhibitory event. In contrast the spontaneous I/E rate ratio had mean 0.43 and median 0.38, indicating far fewer inhibitory events than excitatory events. The median spontaneous I/E rate ratio was lower than the median tone-evoked I/E rate ratio (Wilcoxon sign rank test, p = 3.0 × 10^−7^; Fig. 5A). Tone-evoked excitation is almost always accompanied by tone-evoked inhibition, whereas spontaneous excitation is much less frequently accompanied by inhibition.

**Fig. 5.**
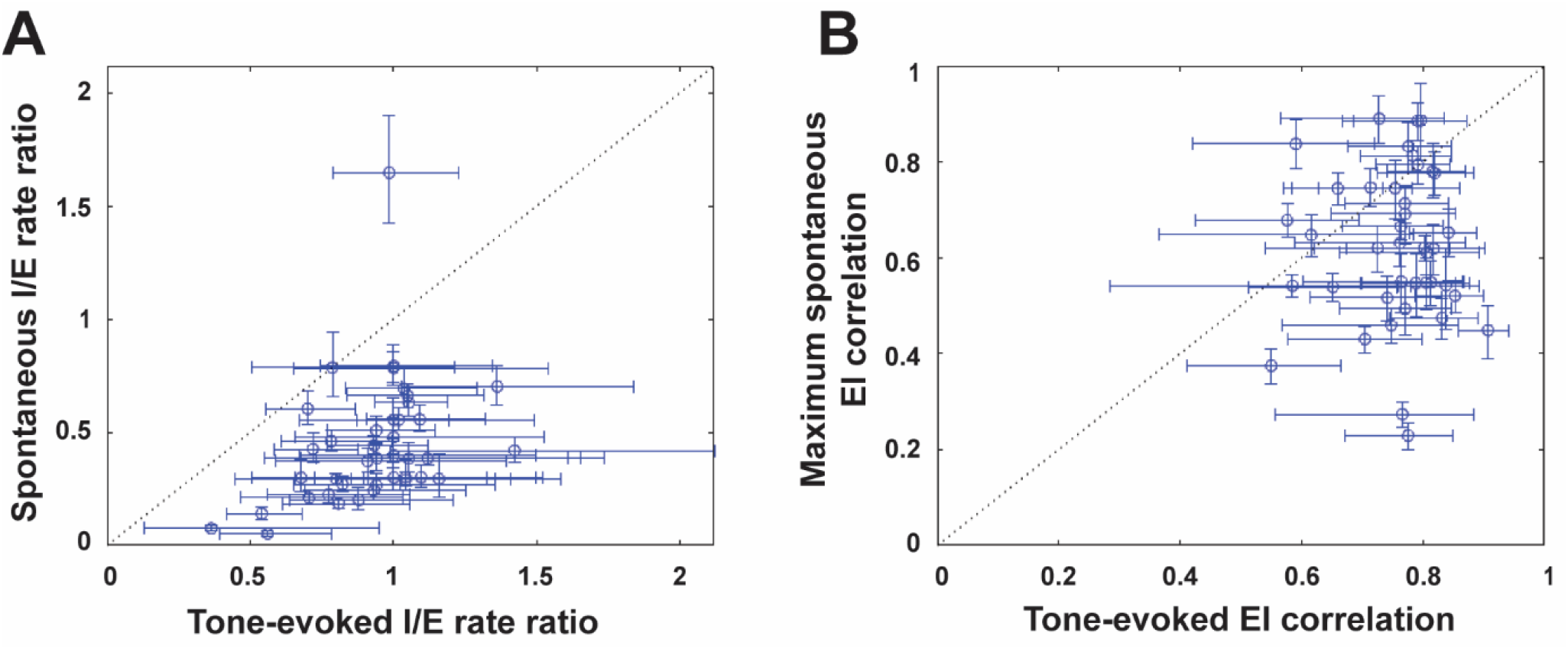
Spontaneous excitatory-inhibitory correlations are lower than tone-evoked excitatory-inhibitory correlations. Data is shown for the 40 neurons in Fig. 4 from which both spontaneous and tone-evoked excitatory and inhibitory currents were recorded (A) Spontaneous I/E rate ratio versus tone-evoked I/E rate ratio. Error bars represent 95% confidence intervals. (B) Maximum spontaneous EI correlation versus tone-evoked EI correlation. Error bars for tone-evoked EI correlations represent 95% confidence intervals; error bars for the maximum spontaneous EI correlations are values corresponding to the 95% confidence limits for spontaneous EI rate ratios.

Next, we compare the maximum spontaneous EI correlation with the tone-evoked EI correlation for each neuron. The maximum spontaneous EI correlation is computed as for Fig. 2B, 4B. The tone-evoked EI correlation is computed as for Fig. 3 (right panels). The median (0.62) maximum spontaneous EI correlation for spontaneous activity was less than the median (0.77) tone-evoked EI correlation (Wilcoxon sign rank test, p = 3.1 × 10^−5^; Fig. 5B). Of the 40 neurons with V-shaped tuning curves, the point estimate for the maximum spontaneous EI correlation was lower than the point estimate for tone-evoked EI correlation in 29 neurons; the 95% upper confidence limit for the maximum spontaneous EI correlation was less than the 95% lower confidence limit for tone-evoked EI correlation in 23 neurons (Fig. 5B).

## Discussion

We have shown that the excitatory event rate is greater than the inhibitory event rate in spontaneous synaptic activity of primary auditory cortex neurons. The ratio by which the spontaneous excitatory rate exceeds the spontaneous inhibitory rate is enough to indicate that the excitatory-inhibitory correlation is lower for spontaneous activity than for tone-evoked responses. The excitatory-inhibitory correlations for spontaneous activity are likely to be lower than our estimated maximum spontaneous EI correlations, as those are upper bounds.

In our analyses we assumed that excitatory and inhibitory synaptic currents can each be reasonably described as a filtered Poisson process, and we only detected spontaneous events with peak amplitude greater than 0.1 of the largest peak. We restricted our analyses those larger events, as the smaller excursions were not so apparently constituted from distinct events, and may have been more affected by voltage-clamp errors producing excitatory contamination in the inhibitory currents. If our assumptions are strongly violated, it is possible for the excitatory-inhibitory correlation to be higher for spontaneous activity than for tone-evoked responses. For example, if in addition to the filtered Poisson process to which the larger events belong, each synaptic current contained a second process that was both highly correlated between excitation and inhibition and had amplitudes smaller than 0.1 of the largest peak, the sum of both processes could have a high excitatory-inhibitory correlation. Visual inspection of our spontaneous activity records suggests that such a second process is unlikely, but better measurements with smaller voltage clamp errors to enable clearer interpretation of the smaller currents will be needed to firmly rule out this possibility.

Our results show that under pentobarbital anesthesia spontaneous excitation and inhibition in the primary auditory cortex and visual cortex are similar (Tan et al., 2013). Our data also show that the balance of excitation and inhibition during spontaneous activity and tone-evoked activity in the primary auditory cortex are different. Hence spontaneous activity does not mainly consist of circuit activation patterns resembling those evoked by random tone sequences. Intriguingly, an analogous differentiation between spontaneous activity and the activity evoked by natural sensory stimuli has been made at the level of population activity in the visual cortex of awake rodents (Stringer et al., 2019). The cortex supports diverse patterns of spontaneous activity, whose features, such as the balance of excitation and inhibition or auto-correlation, depend on behavioral or anesthesia state and cortical area (Haider et al., 2006; Rudolph et al., 2007; Okun and Lampl, 2008; Yaron-Jakoubovitch et al., 2013; Murray et al., 2014). Further work is needed to understand how excitation and inhibition in spontaneous and stimulus-evoked activity change with cortical state and region to provide insight into the circuit mechanisms underlying stimulus encoding (Pachitariu et al., 2015).

## Acknowledgements

This research was supported by a Singapore Ministry of Health National Medical Research Council Open Fund -Individual Research Grant [NMRC/OFIRG/0043/2017] and a National University of Singapore Young Investigator Award [NUSYIA_FY16_P22] to A.Y.Y.T.

